# Activity in the reticulospinal tract scales with handgrip force in humans

**DOI:** 10.1101/2023.10.27.563544

**Authors:** Tyler L. Danielson, Layla Gould, Jason M. DeFreitas, Rob J. MacLennan, Chelsea Ekstrand, Ron Borowsky, Jonathan P. Farthing, Justin W. Andrushko

## Abstract

**Introduction:** The neural pathways that contribute to force production in humans are currently poorly understood, as the relative roles of the corticospinal tract and brainstem pathways, such as the reticulospinal tract (RST), vary substantially across species. Using functional magnetic resonance imaging (fMRI) we aimed to measure activation in the pontine reticular nuclei during different submaximal handgrip contractions to determine the potential role of the RST in force modulation.

**Methods:** Thirteen neurologically intact participants (age: 28 ± 6 years) performed unilateral handgrip contractions at 25%, 50%, 75% of maximum voluntary contraction during brain scans. We quantified the magnitude of RST activation from the contralateral and ipsilateral sides in addition to the contralateral primary motor cortex during each of the three contraction intensities.

**Results:** A repeated measures ANOVA demonstrated a significant main effect of force (*p* = 0.012, η _p_ ^2^ = 0.307) for RST activation, independent of side (i.e., activation increased with force for both contralateral and ipsilateral nuclei). Further analyses of these data involved calculating the linear slope between the magnitude of activation and handgrip force for each ROI at the individual-level. One-sample *t-*tests on the slopes revealed significant group-level scaling for the RST bilaterally, but only the ipsilateral RST remained significant after correcting for multiple comparisons.

**Conclusions:** Here, we show evidence of task dependent activation in the RST pontine nuclei that was positively related to handgrip force. These data build on a growing body of literature that highlights the RST as a functionally relevant motor pathway for force modulation in humans.

**New & Noteworthy:** In this short report, we used a task-based fMRI paradigm to show that activity in the reticulospinal tract, but not the contralateral motor cortex, scales linearly with increasing force during a handgrip task. These findings directly support recently proposed hypotheses that the reticulospinal tract may play an important role in modulating force production in humans.

## Introduction

Within the central nervous system, there are multiple descending pathways that control motor function, and the roles and relative contributions from each pathway may vary substantially across species. In non-mammals, such as reptiles and birds, the main motor pathways descending onto motoneurons originate in the brainstem (1). However, in humans, it is believed that upper limb fine motor control and force modulation is primarily controlled through the corticospinal tract [CST; (2, 3)]; a descending pathway originating from the primary motor cortex (M1). An added difficulty in quantifying the role of the CST is how much its size and functionality differs between species. Within humans, the cortico-motoneuronal connections are substantially more developed than in non-human primates and other mammals (4), suggesting that the role of CST in humans likely cannot be inferred accurately from animal-model research. Increased development of the CST correlates positively with indices of dexterity (5), giving further evidence for the role of the CST in fine, distal motor production. Due to this specialization of the CST within humans, most research studies investigating the mechanisms of human motor control and muscle force have primarily focused on the CST, with equivocal and heterogeneous findings (6, 7).

Recent evidence, however, now suggests the reticulospinal tract (RST); a brainstem pathway thought to be limited in humans to postural and proximal muscle control, may be more involved in upper limb motor control than previously thought (8–10). In support of this, there has been evidence of distal RST innervation in non-human primate models (11) and in humans (8, 9). In addition to its contribution to gross movement control, recent papers have suggested that the RST may also have a role in force modulation (12–14). Some studies have used animal models to study the RST’s role in voluntary force production, but these models require the implantation of electrodes on the RST which is not possible in humans (10, 11, 15–17). For example, Glover and Baker (16) found that the RST appears to influence contraction force in macaque monkeys. Others have examined this control in humans with stroke and spinal cord injury, allowing for models analogous to animal lesion studies (18, 19).

In humans, these studies primarily involve non-invasive stimulation to produce evoked contractions (19–21), while animal models have allowed for studies during voluntary movement (11, 15–17, 22). Examining the role of the RST during voluntary force production is important to elucidate how it contributes to the control of human movement. To our knowledge, no study to date has used functional magnetic resonance imaging (fMRI) to corroborate the role of the RST for controlling force in humans despite the benefits of its non-invasive nature. The purpose of this study was to quantify whether the RST is more active during handgrip contractions at higher compared to lower intensities.

## Materials and Methods

As a secondary analysis on a data set that has previously been published (23) we aimed to determine the relationship between fMRI BOLD signal modulation in the pontine reticular formation nuclei and parametric increases in handgrip force during three separate fMRI runs (Figure 1).

**Figure 1.**
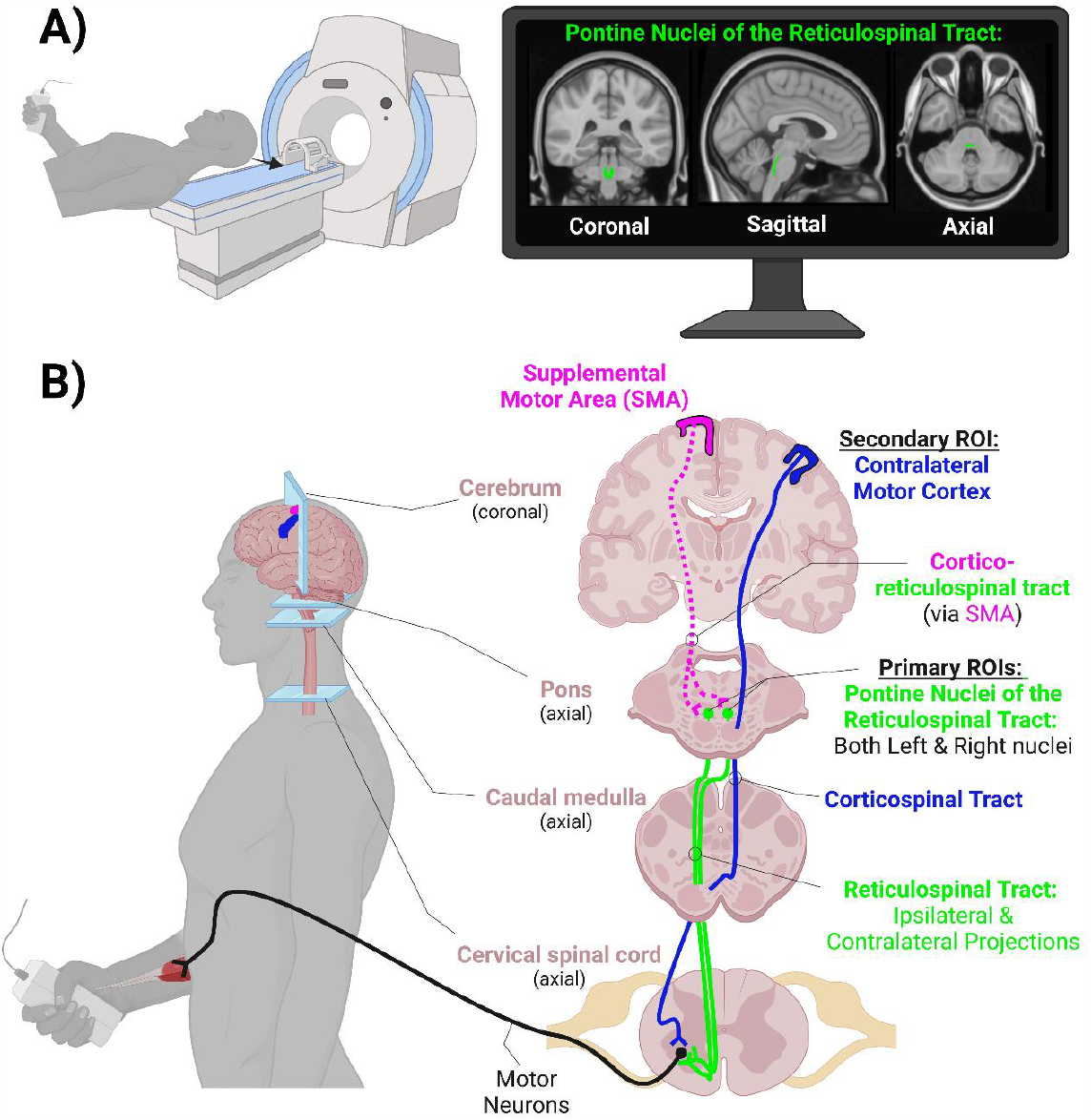
**A)** Handgrip contractions at three different force levels were performed during three separate functional magnetic resonance imaging (fMRI) runs of the cerebrum and brainstem. The monitor in the top right shows our primary region of interest (ROI) in MNI-space, the pontine nuclei of the reticulospinal tract (RST). Section **B)** depicts a simplified diagram of the hypothesized pathways involved in voluntary handgrip force production. For comparison, activity levels in the contralateral motor cortex (M1) were also assessed. Not shown, but also potential contributors, are the contralateral supplemental motor area (SMA) and its bilateral projections to the RST on each side, potential connections to the RST from the premotor cortices, and uncrossed pathways from the ipsilateral M1.

### Participants

Thirteen neurologically intact individuals volunteered to participate in this study (11 right-handed, 2 left-handed; sex: 7 female, 6 male; age: 28 ± 6 years; height: 170.9 ± 9.8 cm; weight: 75.1 ± 16.7 kg). Participants were eligible if they were 18 years or older and were MRI safe. The presence of intracranial metal clips, cardiac pacemaker, or any factor that would prevent a safe MRI scan disqualified individuals to participate.

### Ethical approval

All participants provided written informed consent before participation. The study adhered to the Declaration of Helsinki and received approval from the University of Saskatchewan Biomedical Research Ethics Board (Ethics # Bio 01-125).

### Experimental outline

During a single fMRI session, participants engaged in three experimental tasks. These tasks were executed in a random sequence and consisted of submaximal unimanual right isometric handgrip contractions at 25%, 50%, and 75% of their maximum voluntary contraction (MVC). The session began with the acquisition of a structural MRI brain scan, followed by having the participants perform three MVC’s. To prevent fatigue, a one-minute rest interval was allotted between each MVC attempt. Based on the MVC results, target force levels were set for the three submaximal experimental tasks. Participants then proceeded to complete the three experimental tasks during separate fMRI runs in random order.

### Behavioural motor task

Each of the submaximal handgrip conditions were acquired in separate block design fMRI runs. For each run, participants performed 5 sets × 5 repetitions of either a 25%, 50%, or 75% MVC handgrip visuomotor target matching task with an MRI-compatible hand clench dynamometer (Biopac Systems Inc. Aero Camino Goleta, CA). Task blocks involved 1,650 ms [i.e., corresponding to the repetition time (TR) of the T2* fMRI scan] contractions alternating with 1,650 ms of rest (16,500 ms total task block). Each task block was separated by blocks of complete rest (16,500 ms total rest block). During each fMRI run, the participants wore MRI compatible goggles and viewed a computer screen projection, where they were able to view a custom-built LabView (version 8.6) interface.

Participants saw clear target lines and go/no-go on screen flashing lights and were cued when to contract or relax. The LabView interface was triggered by the MRI through a fibre optic cable that sent a signal at the beginning of each TR. This signal went through a Siemens fMRI trigger converter and passed to the LabView interface via a BNC connection. This allowed for precise continuous synchronization between the MRI and the experimental paradigm computer throughout the scan (24). Target lines were presented relative to the individual’s peak MVC and force feedback was presented as a vertical force bar that was responsive to each participant’s handgrip contraction (i.e., harder contraction resulted in the bar rising vertically). Participants were visually instructed when to contract and when to relax with a green light that would turn on when the participant was to contract. A second red light was programmed to turn on during rest blocks to indicate a sustained rest. Participants were instructed to relax their non-active arm throughout each of the fMRI runs.

### Handgrip motor task processing

The handgrip force data were processed in Matlab using custom scripts (25). Data were processed with a fourth-order 100 Hz low pass Butterworth filter and full-wave rectified. The mean of each contraction was then normalized to the mean Kg-force of the highest MVC and expressed as a percentage of MVC. The mean of each task block (five contractions) was calculated and used in subsequent analyses.

### fMRI parameters

All scans were acquired in a Siemens 3T MAGNETOM Skyra MRI (Siemens Healthcare, Erlangen, Germany). At the start of the session a whole-brain T1-weighted structural scan was acquired using a high-resolution magnetization prepared rapid acquisition gradient echo (MPRAGE) sequence consisting of 192 T1-weighted echo-planar imaging slices (1 mm slice thickness with no gap), with an in-plane resolution of 1 × 1 mm [field of view = 256 × 256; TR = 1,900 ms; echo time (TE) = 2.08 ms].

Each of the experimental fMRI runs, were acquired using T2*-weighted single-shot gradient-echo planar imaging with an interleaved ascending sequence, consisting of 105 volumes (TR = 1,650 ms; TE = 30 ms) of 25 axial slices of 4 mm thickness (1-mm gap) with an in-plane resolution of 2.7 mm × 2.7 mm (field of view = 250) using a flip angle of 90°. The top 2 coil sets (16 channels) of a 20-channel Siemens head coil (Siemens Healthcare) were used. Runs consisted of a 10-volume alternating block design beginning with five volumes for stabilization (task, rest; 105 volumes total).

### fMRI preprocessing

Functional MRI data processing was carried out using FMRI Expert Analysis Tool (FEAT) Version 6.00, as part of FSL [FMRIB’s Software Library; (26)]. Boundary based registration was used to register the functional image to the high-resolution T1-weighted (T1w) image. Registration of the functional images to the T1w image was carried out using FMRIB’s Linear Image Registration Tool [FLIRT: (27, 28)], and then non-linearly (rigid + affine + deformable syn) registered to the MNI152_1mm brain template using antsRegistrationSyN and antsApplyTransforms from Advanced Normalization Tools [ANTs; (29, 30)]. The following pre-processing was applied: motion correction using Motion Correction FMRIB’s Linear Image Registration Tool [MCFLIRT; (28)]; non-brain removal using FNIRT-based brain extraction as implemented in the fsl_anat anatomical processing script (31); spatial smoothing using a Gaussian kernel of FWHM 6 mm; grand-mean intensity normalization of the entire 4D dataset by a single multiplicative factor (32). Next, Independent Component Analysis Automatic Removal of Motion Artifacts (ICA-AROMA) was used remove motion-related noise from each functional run (32). Following ICA-AROMA data clean up, data were high pass temporal filtered with a 0.01 Hz cut off. Time-series statistical analyses were carried out using FMRIB’s Improved Linear Model (FILM) with local autocorrelation correction (33). Z (Gaussianised T/F) statistic images were constructed non-parametrically using Gaussian Random Field theory-based maximum height thresholding with a corrected significance threshold of *p* = 0.05 (34).

### Region of interest signal change

FSL’s featquery tool was used to extract the BOLD signal percent change from three regions of interest (ROI) in participants’ native-space. For each ROI, ANTs were used to transform ROIs from MNI-space to native-space prior to signal change extraction. The percent signal change was extracted from the left contralateral and right ipsilateral Pontine reticular formation nuclei (left: PnO_PnC_l; right: PnO_PnC_r) from the Brainstem navigator toolkit (35). Additionally, the percent signal change was extracted from left contralateral M1 using an ROI mask from the Brainnetome atlas [left M1: A4ul_l; (36)]. The extracted values were then used in subsequent statistical analyses.

### Statistical analysis

An omnibus ROI [contralateral pontine reticular formation, ipsilateral pontine reticular formation] × condition (25%, 50%, 75% MVC) repeated measures Analysis of Variance (RM-ANOVA) was run using the magnitude of brain activation from each of the ROI’s (i.e., BOLD signal percent change from rest) to determine whether RST activation changed with handgrip force modulation. Data were assessed for normality using Shapiro-Wilks in addition to Z_skewness_ and Z_kurtosis_. ANOVA tests are robust against violations to normality (37, 38) and thus minor violations to normality as determined by z_skewness_ and Z_kurtosis_ were considered acceptable for running the parametric model. Sphericity was assessed with Mauchley’s test of sphericity and where violated the Greenhouse-Geisser correction was used.

Hedges *g* effect sizes were calculated and reported to adjust for the small sample size. A Bonferroni adjustment was used to correct for multiple comparisons where appropriate post-hoc testing was conducted. The α-level for significance was set to α = 0.05.

Additionally, for each ROI the slope coefficient of brain activation across the three submaximal handgrip force conditions (percent change of BOLD signal activation / percent target force) was calculated on a subject-by-subject basis using linear regression. The slope coefficients were then analyzed using separate one-sample t-tests to determine if each ROI scales its magnitude of activation across the varying submaximal force outputs. A positive slope indicates activation in that ROI increases with handgrip force. A negative slope would represent a decrease in activation with an increase in handgrip force and a non-significant slope (i.e., not different from zero) would represent that activation did not vary as a function of handgrip force. For this analysis, the α-level was set to α = 0.017 (0.05 / 3 ROIs) and *p*-values ≤ 0.017 were considered significant.

## Results

### Magnitude of BOLD Activation Across Contraction Intensities

RM-ANOVA was performed to test the effect of force (25%, 50%, and 75% MVC) and side (ipsilateral and contralateral) on pontine reticular nuclei activation. Mauchly’s test of sphericity was violated for Force (*p* = 0.023) and Force × Side (*p* = 0.006). There was a significant main effect for force [*F*(1.337, 16.039) = 5.306, *p* = 0.012, η_p_^2^ = 0.307], but no main effect for side [*F*(1,12) = 0.413, *p* = 0.533, η_p_^2^ = 0.033]. The interaction effect (Force × Side) was also not significant [*F*(1.243, 14.921) = 0.957, *p* = 0.364, η_p_^2^ = 0.074]. A Bonferroni post hoc analysis showed that activation of pontine nuclei at 75% MVC was significantly greater than at 25% MVC (MD = -0.2, SE = 0.068, *p* = 0.02) as well as at 50% MVC (MD = -0.179, SE = 0.068, *p* = 0.042). The difference in activation between 25% and 50% MVC was not significant (MD = -0.021, SE = 0.068, *p* = 1.000; Figure 2A).

**Figure 2.**
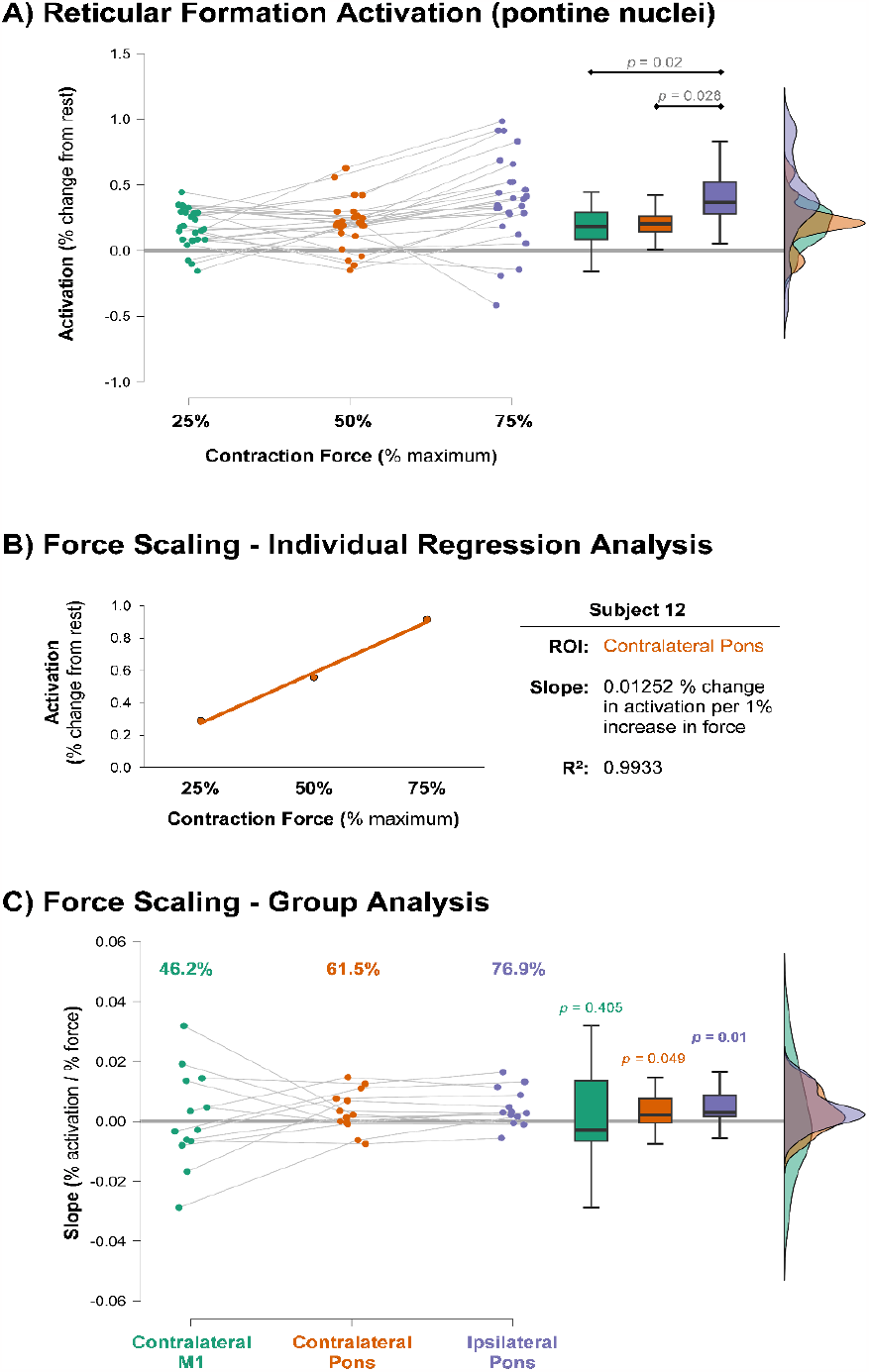
The top panel **(A)** shows the activation in the pontine nuclei of the reticular formation during contractions of varying intensities. The contralateral and ipsilateral nuclei are collapsed together because there was neither a side × force intensity interaction nor a main effect for side as a factor. Pontine activation did show a main effect for force intensity, with post-hoc pairwise comparisons shown above the box plots. The middle panel **(B)** shows the relationship between % activation and % force from a single region of interest (ROI) from a representative subject. The relationship was fit with a linear model, and the slope interpreted as the degree to which activation for that region changes with increases in force. Panel **(C)** shows the slope coefficients from each subject for each ROI. The values above the scatter plot demonstrate the percent of subjects that exhibited a positive slope (i.e., positive scaling in activation with increasing force) for each ROI. The *p*-values above the box plots are from one-sample *t*-tests, showing that the mean of the slopes was significantly different than zero in the pons before correcting for multiple comparisons, but not in M1. Only the ipsilateral pontine reticular nuclei significantly scaled after statistical correction (corrected α = 0.017). The thick gray horizontal lines in A and C extend the zero point for easy comparison.

### ROI BOLD Activation Scaling with Force

Linear regression analyses were conducted on each ROI for each subject to determine the slope coefficient, which quantifies the rate of increase in activation with increasing contraction intensity (Figure 2B). Next, using separate one sample *t*-tests of the individual slopes, we showed at the group-level pontine reticular activation was significantly different from zero on both the ipsilateral [*t*(12) = 2.672, *p* = 0.01, *g* = 0.712] and contralateral [*t*(12) = 1.792, *p* = 0.049, *g* = 0.478] sides before correction; however, after a manual Bonferroni correction for multiple comparisons, only the ipsilateral side remained significant. The contralateral M1 was not statistically different from zero [*t*(12) = 0.245, *p* = 0.405, *g* = 0.065], indicating it did not reliably scale with force (Figure 2C).

## Discussion

The present study examined the relationship between unimanual handgrip force production and BOLD signal activation of the contralateral and ipsilateral pontine reticular nuclei of the reticulospinal tract, as well as the contralateral M1 for comparison. Our primary finding is that activation within the RST consistently scaled linearly with increased force production, which was not the case for the contralateral M1. The traditional understanding is that the RST primarily innervates axial and proximal muscle groups in humans (39). However, over the past 15 years a small body of literature has begun to challenge this belief (8, 9, 11, 40) with research suggesting the RST may reach distal musculature of the arms and hands and modulate force (12–14). With this task, we have shown, for the first time, pontine reticular nuclei BOLD signal activation significantly scales with handgrip force, where a positive relationship between the magnitude of activation and force output are observed.

Specifically, we observe significant scaling in the ipsilateral pons (with a large effect size) after correcting for multiple comparisons. There was also scaling with a moderate effect size in the contralateral pons. Additionally, the repeated measures ANOVA showed that side (ipsilateral vs. contralateral) had no effect on activation across the force conditions. These findings, paired with the lack of scaling in contralateral M1 provide some support for the notion of the RST pathway contributing to high force outputs, rather than the traditional belief that human distal motor control is exclusively governed by contralateral cortical regions. Schmidt et al. (41) showed that activity in neurons from M1 do not change with increases in right hand force in non-human primates (Rhesus macaque monkeys). They concluded that M1 was responsible for activating a muscle but not modulating force once it was active. Our data in humans, support their conclusion, as we found no robust support for the role of M1 in force modulation, and add to it with evidence that force modulation may be driven by the RST.

During isometric contractions, the recruitment of motor units tend to occur in an orderly fashion from low-threshold to higher-threshold (42). As force demand increases, larger and higher-threshold motor units are recruited (43–46), and the firing rates of smaller, lower-threshold motor units that were already active increase. These recruitment properties may also affect neuromuscular plasticity, as the repeated activation of the larger motor units with high-intensity resistance training often lead to greater gains in strength compared to lower-intensity training (47). In addition to the impact of high-intensity contractions on motoneuron recruitment, and strength, a recent review by Hortobágyi et al. (48) reported high-intensity resistance training also produces greater motor-related neuroplasticity in the neurologically intact young adult brain. However, mechanistic studies into the specific neural pathways and adaptations to resistance training that affords this motoneuronal recruitment remains incompletely understood. This incomplete mechanistic understanding is due in part to research focussing on the CST, which has resulted in equivocal insight. Here, we add to the growing body of literature that pinpoints the RST as a relevant pathway for force control with a handgrip task, which, until recently (12–14), was thought to be exclusively controlled by the CST (49). Specifically, we show evidence for intensity dependent activation of the RST that is correlated with force output, which suggests this is a task relevant pathway for handgrip force. Further, based on the principles of use-dependent plasticity, the repeated preferential activation of the RST with high-intensity contractions over the course of a resistance training intervention may lead to the strengthening of this pathway’s integrity in a manner that facilitates strength adaptations. Recent findings have led to proposals that the RST may have a role in modulating motor unit discharge rates, which would in turn alter force and rate of force production (14). Škarabot et al. (14) proposed hypothesis about the role of RST in modulating force is supported by the findings of this study.

Evaluating the RST in applied settings presents its own set of challenges, such as the need for external stimuli (e.g., Transcranial Magnetic Stimulation or cervical medullary electrical stimulation) and perturbations (e.g., loud acoustic startle sounds ≥ 110 dB), in addition to a high variance in responses. Here, using fMRI, we show a non-invasive evaluation of the RST in a controlled environment during voluntary motor behaviours. We argue this is an important contribution to the field investigating the role of the RST, as these data provide support to previous findings without the need for external perturbations. Further, when paired with previous research (12, 13, 15), our findings indicate that the RST may be a functionally relevant pathway contributing to muscle force and, by extension, possibly chronic strength training adaptations.

### Limitations and future directions

Investigating patterns of brain activation during high-intensity contractions with fMRI is not a trivial task, as head motion commonly occurs during effortful movements, and can result in the loss of data due to motion related contamination. With that, the investigation of fMRI related BOLD signal activation during different types of motor tasks may not be possible, and thus these experiments are limited to simpler isolated hand movement patterns that are less likely to produce high amounts of motion.

Though we present evidence for the role of the RST in gross distal hand movement, future research may consider testing RST activation during finer distal movements. By focusing on dexterous movements such as flexion, extension, abduction, adduction, and circumduction, RST activation bias toward certain types of movement may be determined. Future research into this area should investigate the contributions of the RST to different tasks and their adaptations to training interventions with varying manipulations. A deeper understanding of how the RST contributes to motor function could lead to better rehabilitative modalities for those with dysfunction of the CST, such as individuals with stroke (8, 12). Additionally, we focused only on the pontine nuclei of the RST. The medullary nuclei had to be excluded because the medulla was cut off from the scan for many of the participants in favour of capturing complete cortex. Future work should examine whether the medullary nuclei also play a potential role in force modulation. Finally, future research would benefit from replication of this experiment but with a larger sample size to build confidence in data interpretation and robustness of findings.

## Conclusions

In a small cohort of neurologically intact young adults, we demonstrate that activation of the pontine reticular nuclei of the reticulospinal tract scales in magnitude with handgrip force. These data provide insight into the potential role of the RST during distal muscular force modulation, a concept most often considered under exclusive control by the CST in humans. These data add to the growing body of literature suggesting a potentially important role of the RST for neuromuscular strength adaptations in humans and lays the foundation for which future research can further investigate RST function to disentangle the neural underpinnings of force modulation and the neural adaptations governing increases in strength.

## Data availability Statement

Data are available upon reasonable request

## Acknowledgements

We would like to thank Doug Renshaw, Sul Ross State University, for this assistance with data collection, and Shawn Reinink, University of Saskatchewan, for his contribution to the LabView software interface using in this experiment.

## Grants

This study was funded by a Natural Sciences and Engineering Research Council of Canada (NSERC) Discovery Grant: 2016-0529, awarded to Jonathan P. Farthing, in addition to an NSERC Discovery Grant: 183968-2013-25, awarded to Ron Borowsky.

## Disclosures

The authors have none to declare

## Author Contributions

J.W.A. and L.G. collected the data. J.W.A. processed the data and assisted with statistical analyses and the writing of the manuscript. T.L.D. took the lead on writing the manuscript and performed the statistical analyses. J.M.D. conceived the research questions for which these data were used to answer, created figures, contributed to writing, edited and approved the final version of the manuscript. R.L.M. assisted with writing the manuscript and approved the final version. C.E. contributed to data processing and analysis. J.P.F. and R.B. were involved in obtaining the grant funds and ethical approvals for the larger study of which the data from this study are from, they were also involved in the original study design, editing and approving the final version of the manuscript.

## Notes

### Competing Interest Statement

The authors have declared no competing interest.

### Summary of Updates

Adjusted wording to statement in discussion and added an individual to acknowledgements for their prior contribution

